# SEGMENTATION OF DYNAMIC TOTAL-BODY [^18^F]-FDG PET IMAGES USING UNSUPERVISED CLUSTERING

**DOI:** 10.1101/2023.06.20.545535

**Authors:** Maria K. Jaakkola, Maria Rantala, Anna Jalo, Teemu Saari, Jaakko Hentilä, Jatta S. Helin, Tuuli A. Nissinen, Olli Eskola, Johan Rajander, Kirsi A. Virtanen, Jarna C. Hannukainen, Francisco López-Picón, Riku Klén

## Abstract

Clustering time activity curves of PET images has been used to separate clinically relevant areas of the brain or tumours. However, PET image segmentation in multi-organ level is much less studied due to the available total-body data being limited to animal studies. Now the new PET scanners providing the opportunity to acquire total-body PET scans also from humans are becoming more common, which opens plenty of new clinically interesting opportunities. Therefore, organ level segmentation of PET images has important applications, yet it lacks sufficient research. In this proof of concept study, we evaluate if the previously used segmentation approaches are suitable for segmenting dynamic human total-body PET images in organ level. Our focus is on general-purpose unsupervised methods that are independent of external data and can be used for all tracers, organisms, and health conditions. Additional anatomical image modalities, such as CT or MRI, are not used, but the segmentation is done purely based on the dynamic PET images. The tested methods are commonly used building blocks of the more sophisticated methods rather than final methods as such, and our goal is to evaluate if these basic tools are suited for the arising human total-body PET image segmentation. First we excluded methods that were computationally too demanding for the large datasets from human total-body PET scanners. This criteria filtered out most of the commonly used approaches, leaving only two clustering methods, k-means and Gaussian mixture model (GMM), for further analyses. We combined k-means with two different pre-processings, namely principal component analysis (PCA) and independent component analysis (ICA). Then we selected a suitable number of clusters using 10 images. Finally, we tested how well the usable approaches segment the remaining PET images in organ level, highlight the best approaches together with their limitations, and discuss how further research could tackle the observed shortcomings. In this study, we utilised 40 total-body [^18^F]fluorodeoxyglucose PET images of rats to mimic the coming large human PET images and a few actual human total-body images to ensure that our conclusions from the rat data generalise to the human data. Our results show that ICA combined with k-means has weaker performance than the other two computationally usable approaches and that certain organs are easier to segment than others. While GMM performed sufficiently, it was by far the slowest one among the tested approaches, making k-means combined with PCA the most promising candidate for further development. However, even with the best methods the mean Jaccard index was slightly below 0.5 for the easiest tested organ and below 0.2 for the most challenging organ. Thus, we conclude that there is a lack of accurate and computationally light general-purpose segmentation method that can analyse dynamic total-body PET images.

**Key points:**

- Majority of the considered clustering methods were computationally too intense even for our total-body rat images. The coming total-body human images are 10-fold bigger.
- Heterogeneous VOIs like brain require more sophisticated segmentation method than the basic clustering tested here.
- PCA combined with k-means had the best balance between performance and running speed among the tested methods, but without further preprocessing, it is not accurate enough for practical applications.

**Funding:** Research of both first authors was supported by donation funds of Faculty of Medicine at University of Turku. JCH reports funding from The Academy of Finland (decision 317332), the Finnish Cultural Foundation, the Finnish Cultural Foundation Varsinais-Suomi Regional Fund, the Diabetes Research Foundation of Finland, and State Research Funding/Hospital District of Southwest Finland. KAV report funding from The Academy of Finland (decision 343410), Sigrid Juselius Foundation and State Research Funding/Hospital District of Southwest Finland. JH reports funding from The Finnish Cultural Foundation Varsinais-Suomi Regional Fund. These funding sources do not present any conflict of interest.

**Data availability:** The codes used in this study are available from Github page https://github.com/rklen/Dynamic_FDG_PET_clustering. The example data used in this study have not been published at the time of writing.

## 1. Introduction

Dynamic positron emission tomography (PET) studies provide a series of threedimensional (3D) images over time. In PET imaging, tracer with radioactive component is usually injected into the body and the spreading of the tracer is followed by recording the gamma radiation resulting from the radioactive decay. Therefore, the dynamic PET data comprises of tracer radioactivity over time in each 3D pixel called voxel. The vectors containing the tracer activity over time are called time-activity curves (TACs) and each voxel has its own TAC. To study the activity of a specific tissue, the images are segmented and one or more volumes of interest (VOIs) are taken into further analysis. A VOI is a 3D version of the commonly used term ‘region of interest’ (ROI). Traditionally, the VOIs have been manually segmented based on additional magnetic resonance imaging (MRI) or computed tomography (CT) scan, but those imaging protocols provide structural information rather than regions with distinct distribution of the used PET tracer. Therefore, structurally similar, but functionally different areas will be ignored in MRI or CT based segmentation, which might lose clinically important information. In addition, manual segmentation is user-dependent and time consuming [1]. An automatic segmentation method for PET images could avoid some of these problems and make PET image analysis more efficient and accurate.

Many automatic segmentation methods have been introduced, tumour segmentation being the single most common application. Due to the small size of the traditional PET scanners, typically the automatic segmentation focuses on a small area of a body, such as the brain. Section Literature review’ provides an overview of the proposed methods and shows that there are several unsupervised methods evaluated with small body-area only, but majourity of the new segmentation methods are different machine learning approaches. The main problems with the published methodology are that 1) most of the methods are designed for a specific purpose or tracer, 2) implementations are typically not publicly available, and 3) many combine PET images with some additional information. While the machine learning based approaches are computationally light to use after the training phase, their usage is limited to images somewhat similar than the utilised training data, and implementations of the finalised methods are usually not available. General-purpose unsupervised methods instead are typically computationally heavier for the end-user and the sophisticated fine-tuned methods suffer from the same lack of available implementation than the unsupervised methods.

New PET scanners, which enable human total-body pet scans, have become more and more popular and they have a great potential for novel scientific discoveries and medical applications. Thus, motivation to segment very large PET images has suddenly exploded, yet the previous studies introducing unsupervised automatic segmentation methods have mainly focused on a single organ or on a very limited area of the body. While implementations of the ready methods are often not available, many of them utilise classical clustering algorithms with different pre- and post-processings. The aim of this study was to test if previously used clustering methods can be used to segment large dynamic total-body PET images. A clustering method needs to a) be computationally light and b) provide biologically meaningful segments in order to be useful tool for automatic segmentation of large modern PET images. The first criteria is important as the modern total-body scans generate tens of millions of TACs per image and the number is likely to further increase as the scanners become more and more accurate and comprehensive.

According to our tests, only two clustering approaches, k-means and Gaussian mixture models (GMM), were fast and light enough to analyse the modern images within reasonable time. Thus we had to exclude some previously successfully used clustering methods, such as spectral clustering, from further analyses. We tested these two clustering methods’ capability to identify VOIs representing heart, brain, and kidneys defined from each analysed PET image by comparing the clustering results to the manually segmented organs using Jaccard index as an evaluation metric. Two different preprocessing approaches, namely principal component analysis (PCA) and independent component analysis (ICA), were tested and we also briefly investigated if the defined organ specific VOIs consist of multiple clusters.

This study consists of sections Literature review, Materials and methods, Results, and Discussion and conclusions. In the Literature review, we introduce different published approaches to automatically segment PET images. The summarised methodology varies on the target application, intended input data, and theoretical principles. In section Materials and methods we provide details important for the reproducibility of this study, including description of the used datasets, summary of the tested methods, and explanation of the followed test design. The Results section contains the outcomes of the tests and evaluations described in subsection test design. There we report the number of clusters that provided the best segmentation results together with the Jaccard indices indicating how well the tested methods performed in different datasets. Also some biologically interesting observations of the obtained automatic segmentations are presented in the Results section. Finally, in the section Discussion and conclusions, we summarise our key observations and discuss the limitations of this study and the available methodology, as well as present ideas how the automatic segmentation could be improved.

## Literature review

Here we provide a summary of the proposed methods to segment PET images. We focus on unsupervised methods not requiring other image modalities, but also provide a bried introduction of other types of approaches. Notably, machine learning (ML) approaches requiring training data are not considered as unsupervised despite them appearing as such for the end-users. Many of the methods and pipelines introduced below have reported impressive results, but do not provide an implementation so that other researchers could apply the proposed method on their own applications without coding it from a scratch. This issue is common to all types of approaches introduced here. Background filtering is sometimes referred to as segmentation ([2]), but in this study our main focus in on segmenting multiple areas instead of binary case. However, tumour segmentation is the most common application of binary segmentation and older approaches suggested for it have been compared in the literature [3, 4]. Also, several modern deep learning methods have been developped for segmenting tumours, as briefly introduced here.

One of the possible ways to automatically segment PET images in unsupervised manner is to cluster TACs, and this technique with multiple different variations has been utilised in previous studies. For instance, Wong *et al*. [5] proposed a method similar to k-means clustering, which was based on a weighted least-square distance, whereas Vogel *et al*. [6] utilised BASC-clustering combining bootstrapping and k-means with sophisticated preand post-processing. K-means is utilised also in the approach by Kim *et al*. as they combine it with region growing [7]. In addition, spectral clustering has been utilised in several studies as Mouysset *et al*. [8] segmented dynamic PET images with kinetic spectral clustering method and Zbib *et al*. [9] suggested using spectral clustering with automatically inferred parameters. Several other classical clustering approaches have been applied as well. For example, Chen *et al*. [10] used an expectation maximisation based clustering algorithm with Markov Random Field models, Guo *et al*. proposed an approach combining fast pre-clustering with hierarchical clustering [11] and later expanded their approach to better suit dynamic 3D PET images [12], and Ashburner *et al*. [13] introduced a modified mixture model algorithm, which clusters based on the shapes of the TACs rather than their absolute scaling. In addition, Parker and Feng [14] used a segmentation algorithm based on Mumford-Shah energy minimisation and pre-processed the data with PCA. Also Kimura *et al*. used PCA as preprosessing prior to their modelbased clustering [15]. In a relatively new study by Cui *et al*., the authors breafly compare different approaches and recommend using definition density peak clustering combined with pre-screening and denoising to segment tumours from PET images [16]. The role of the preprocessing is also highlighted in the iterative affinity propagation based approach by Xu *et al*. [17]. While clustering is the most common approach for this task on dynamic images, also contouring [18, 19] and level set analysis [20] has been applied. Table 1 summarises what type of PET images different unsupervised segmentation approaches introduced in the literature are designed for.

**Table 1.** Summary of target applications of different unsupervised segmentation methods for PET images. Column ‘Studied area’ indicates the body part segmented in the reference in the last column. Columns ‘Dynamic’, ‘Dimensions’, and ‘Organism’ state if the method was designed for dynamic data including multiple time points, number of spatial dimensions of the images to be segmented, and the specie used as an example in the reference, respectively. Due to the lack of access to the full studies [20, 15], some information is missing for them. Notably, the dimensions and dynamic/static nature indicated here refers to the final input for the segmentation as used in the reference. It may differ from the original raw images as e.g. dynamic images may be summed over time points prior to segmentation to mimic a static image or 3D image can be sliced into 2D images.

| Studied area | Dynamic | Dimensions | Organism | Reference |
| --- | --- | --- | --- | --- |
| multi-organ phantom, real brain, real lung | yes | 2D | phantom and human | [5] |
| brain | no | 3D | human | [6] |
| brain (real and phantom) | yes | 3D | phantom and human | [7] |
| brain (real and phantom) | yes | 2D | phantom and rat | [8] |
| brain (real and phantom) | yes | 2D | phantom and rat | [9] |
| simulated, brain | yes | 2D | artificial and human | [10] |
| brain | yes | 2D and 3D | human | [11] |
| brain | yes | 3D | human | [12] |
| brain | yes | 2D | human | [13] |
| brain (real and phantom) | yes | 3D | phantom | [14] |
| brain | yes | - | human | [15] |
| lung, breast | no | 2D | human | [16] |
| several | no | 3D | phantom, human, and rabbit | [17] |
| total-body (real and phantom) | yes | 3D | phantom and rat | [18] |
| prostate | yes | 3D | human | [19] |
| - | yes | 3D | mouse | [20] |

PET images are very heterogeneous based on the used tracer, which makes them nonideal image modality for supervised machine learning despite their advantages related to functional information. Yousefiri *et al*. provide a good summary of the strengths and weaknesses of PET imaging based machine learning [21]. Due to the issues with PET images, including heterogeneity, low resolution, and often poor visibility of anatomical structures, PET images are often paired with CT in different ML approaches. While segmentation of different tumours is the most popular application of ML-based PET/CT segmentation [22, 23, 24, 25, 26], for example Scarinci *et al*. and Sundar *et al*. have introduced more general total-body segmentation methods combining these two image types [27, 28]. While majority of the ML-segmentations use some other image modality besides PET, also purely PET-based ML segmentation has been proposed for tumour detection [21, 29, 30], for more general abdominal area segmentation [31], and even to segment multiple organs from total-body PET images [32]. Taghanaki *et al*. also compare their approach to popular ML-based segmentation tools not designed specifically for PET images [32].

Notably, currently active research focuses mostly on training data based ML-approaches. While they can be very strong tools with excellent performance, their usage is always somewhat limited by the utilised training data, and we believe that both unsupervised and ML-approaches should be actively developed. Besides unsupervised methods for PET images only and modern supervised ML-approaches, there are few segmentation methods not suiting either cathegory. For example, Saad *et al*. introduced a semi-supervised segmentation appraoch based on uncertainty principles [33], which Mirzaalian *et al*. later applied to do the segmentation parallel to motion correction [34]. Also some approaches for unsupervised tumour segmentation using both PET and CT or MRI simultaneusly have been introduced in the literature [35, 36, 37].

## 3. Materials and methods

### 3.1. Data

In this study, we used total-body dynamic 3D PET images of 40 male Sprague-Dawley rats (mean weight 487 grams with standard deviation of 90 grams), which were scanned at Turku PET Centre using Inveon MM Platform by Siemens Molecular Imaging. The scans contained 50 time frames (30 · 10s, 15 · 60s, 5 · 300s). The used tracer was [^18^F]fluorodeoxyglucose ([^18^F]-FDG) and the average dose of it was 20.8 MBq with standard deviation of 1.19 MBq. At the time of imaging, the rats were 8-35 weeks old, under different diets and exercise programs (but fasting for the last 4 hours), and they were in insulin clamp. The images contained 128 · 128 · 159 = 2605056 voxels making them approximately ten-fold smaller than the human total-body scans. Notably, all the voxels were not clustered. Those voxels with mean radioactivity over time below the average mean activity over all voxels were defined as background and were not used for clustering analysis. Thus on average 20.4% of voxels were used in clustering (Supplementary Figure 1). Detailed listing of numbers and proportions of voxels used for clustering is available at https://github.com/rklen/Dynamic_FDG_PET_clustering. Because the images were clustered independently from each other, the activity values in TACs were used as such without further scaling.

In addition to the rat images, we clustered three dynamic total-body PET images of humans (1 female and 2 male). The used tracer was the same [^18^F]-FDG as in the rat data. All human subjects were healthy, as determined by routine laboratory tests, oral glucose tolerance test, and medical examination prior to inclusion. The weights of the three individuals were 79.9 kg, 70.2 kg, and 64.3 kg, and the injected tracer doses were 100.71 MBq, 106.55 MBq and 106.93 MBq, respectively. The scan was started immediately after bolus injection of [^18^F]-FDG. The scans had 13 time frames (1 · 60s, 6 · 30s, 1 · 60s, 3 · 300s, 2 · 600s). These images contained 440 · 440 · 354 = 68534400 voxels, and 8.7-10.0 % of them were used for clustering. The selection criteria for the voxels to be clustered was the same mean-based threshold than with the rat images. The reference number of the ethical committee decision related to the human data is 14/1801/2022 (Hospital District of South-Western Finland).

### 3.2. Clustering methods

Multiple interesting segmentation approaches were considered for this study, but eventually excluded due to too long running time or memory consumption for practical applications on modern big images. The excluded clustering methods include spectral clustering, agglomerative clustering (i.e. hierarchical clustering), affinity propagation, HDBSCAN, and linkage vector from Python package fastcluster. Besides different clustering methods, also other types of segmentation approaches were considered, but excluded due to computational reasons. Such excluded methods include slic, watershed, random walker, morphological geodesic active contouring, and morphological active contours without edges. Only two clustering methods, k-means and Gaussian mixture model, were usable with our test images.

#### K-means clustering with principal component analysis (PCA)

Principal component analysis (PCA) is a popular unsupervised multivariate analysis technique, which can reduce high dimensional data into a smaller number of components. These components are called Principal Components and they explain most of the variance in the original multivariate data with the least loss of information. PCA uses a transformation matrix formed from the eigenvectors of the covariance matrix of the vector data to decorrelate the data. In this study, the PCA was performed on the TAC data with the Python function PCA, which implements the probabilistic PCA model of Tipping and Bishop [38]. Because we dealt with a large number of voxels, the singular value decomposition (SVD) solver in function PCA runs a randomised SVD by the method of Halko *et al*. [39] and Martinsson *et al*. [40].

#### K-means clustering with independent component analysis (ICA)

Independent component analysis (ICA) is a method that can be used to find a linear representation of non-Gaussian data so that the components are statistically as independent as possible. This representation can capture the essential structure of the data in many applications, such as feature extraction and signal separation. In this study, ICA was performed on the TAC data with the Python function FastICA, whose implementation bases on Hyvärinen *et al*. [41].

K-means is a method that aims to separate *n* samples into *k* clusters based on the distance between the samples and the initialised cluster centroids. The k-means implementation used in this study was the Python function k-means, which minimises within-cluster sum-of-squares called inertia:

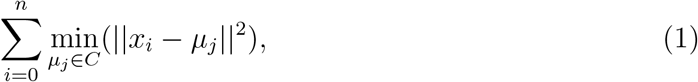

where *x*_*i*_ is the *i*th sample and *μ*_*j*_ is the *j*th cluster centroid from cluster *C*. K-means assigns each sample to the closest centroid using the inertia criterion (1). After that, k-means calculates new centroids by taking the mean value of all the samples in each cluster. This process is repeated until the difference between the old and the new centroids is less than a threshold. K-means clustering uses random centroid initialisation, thus repeated replications may yield different results. An additional drawback of k-means is that the number of clusters must be defined in advance. PCA and ICA can be combined with k-means (or any other clustering approach) by preprocessing the data with either of them prior to the actual clustering. Such preprocessing reduces the dimension of each data point to be clustered to the chosen number of independent or principal components.

#### Gaussian mixture model (GMM)

The third tested method is a probabilistic model called Gaussian mixture model (GMM). Mixture models generally assume that any distribution can be conveyed as a mixture of distributions of some known parametrisation. In the case of GMM, the data is assumed to have been generated by more than one Gaussian distributions. Clustering happens by assigning a label to the Gaussian that contributes the largest probability. A GMM is solved using Expectation-maximisation (EM) algorithm, which calculates maximum likelihood estimates of the parameters of each component distribution. The EM algorithm is guaranteed to converge, so a locally optimal solution is always obtained. The GMM algorithm implementation used in this study was a Python function GaussianMixture. All the utilised clustering related functions were from Python library sklearn associated with version 1.1.2 of scikit-learn, and the used Python version was 3.10.

### 3.3. Test design

Figure 1 summarises the test design of this study. In order to validate the clustering results, we needed known VOIs for heart, brain, and kidneys from each image. Those were obtained from the 40 PET images utilised in this study by manually drawing the organs under the supervision of a biologist with over 15 years of experience on rat models and PET image analysis. The VOIs were drawn with Carimas software (version 2.10) [42]. Importantly, the gold standard to define VOIs is to obtain them directly from the PET images based on tracer activity instead of corresponding CT images visualising static structure. Thus we only used PET images to obtain VOIs. The rounded average numbers of voxels with their standard deviations (in parenthesis) of heart, brain and kidneys were 2834 (757), 4846 (888), and 3054 (1634), respectively. The corresponding mean volumes were 1359, 2325, and 1465 *mm*^3^. The average TACs of these VOIs are visualised in Supplementary Figure 2. Notably, the kidneys did not always fully fit into the scanned area, which explains the high variation in their sizes.

**Figure 1.**
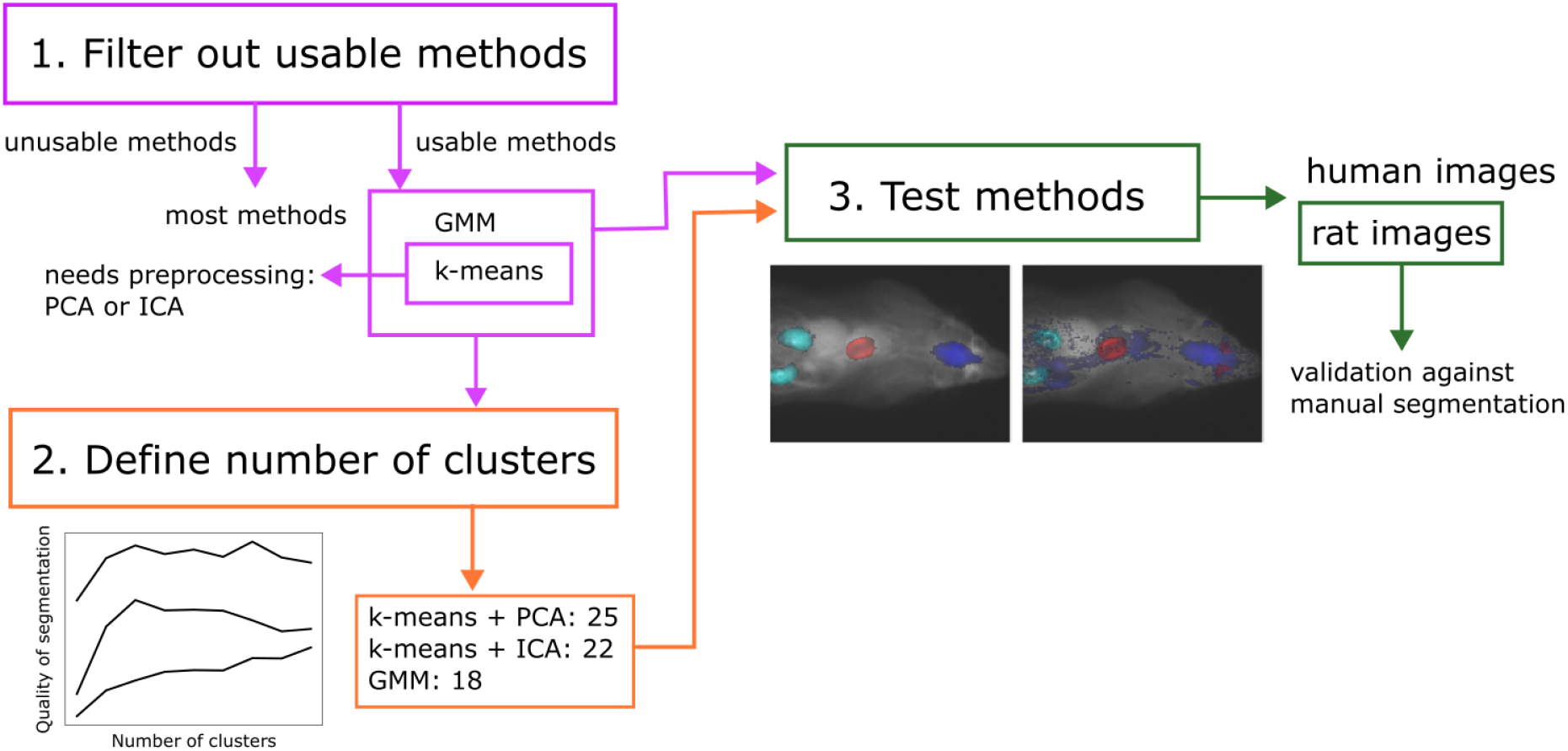
Workflow of the study

The performance of each method for each organ was evaluated using Jaccard index, which describes how well the cluster and the manual segmentation of the analysed VOI overlap. Jaccard index 1 indicates perfect match between the cluster and the manually drawn segment, whereas Jaccard index becomes 0 if the VOIs do not overlap at all. Jaccard index is formally defined as the rate of the intersection and the union of the two VOIs:

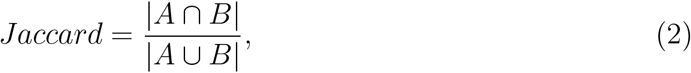

where *A* is the set of voxels belonging to the cluster representing the analysed organ, and *B* is the set of voxels manually segmented to form the VOI for the organ. Notations *A B* and *A B* indicate the number of voxels in the intersection and union of *A* and *B*, respectively. For each organ, the cluster with the highest Jaccard index was selected to represent it. The statistical significance between the Jaccard indices from two methods was calculated using two-tailed Wilcoxon signed-rank test with the conventional p-value cutoff of 0.05 for significance.

As all of the tested methods require a number of clusters to use, we tested cluster numbers 5, 10, 15, …, 45 for all methods with the 10 images, which were acquired first. Then, for each method separately, we selected the average of the best (defined as the highest Jaccard index) organ specific cluster numbers for the further analyses with the remaining 30 images. For other parameters we used the implementations’ default values. For k-means clustering this means that the initial cluster centroids were selected using sampling based on an empirical probability distribution of the points’ contribution to the overall inertia, and the maximum number of iterations was set to 300. The most important default parameters for GMM were covariance type, which is defined so that each component has its own general covariance matrix, the number of EM iterations to perform (set to 100), and parameter init params, which defines how to obtain the initial weights, means, and precisions (set to ‘kmeans’). The remaining parameter defaults can be found from python package sckikit-learn’s instruction pages at https://scikit-learn.org/stable/modules/generated/sklearn.cluster.KMeans.html and https://scikit-learn.org/stable/modules/generated/sklearn.mixture.GaussianMixture.html.

A computer with 16GB of RAM and Intel Pentium Gold processor G6405T (CPU 3.50GHz) was used in the analyses and the reported running times are obtained with it.

## 4. Results

### 4.1. Number of clusters

First, the number of clusters for each method was defined. The tested cluster numbers were between 5 and 45, and 10 images were analysed. Our results show (Figure 2) that PCA had the best performance with the largest number of clusters and GMM’s performance was at its best with the smallest number of clusters. For further analyses, we used 25 clusters for PCA, 22 for ICA, and 18 for GMM. Notably, GMM produced errors with some analyses, particularly with large cluster numbers, so those results are missing from Figure 2.

**Figure 2.**
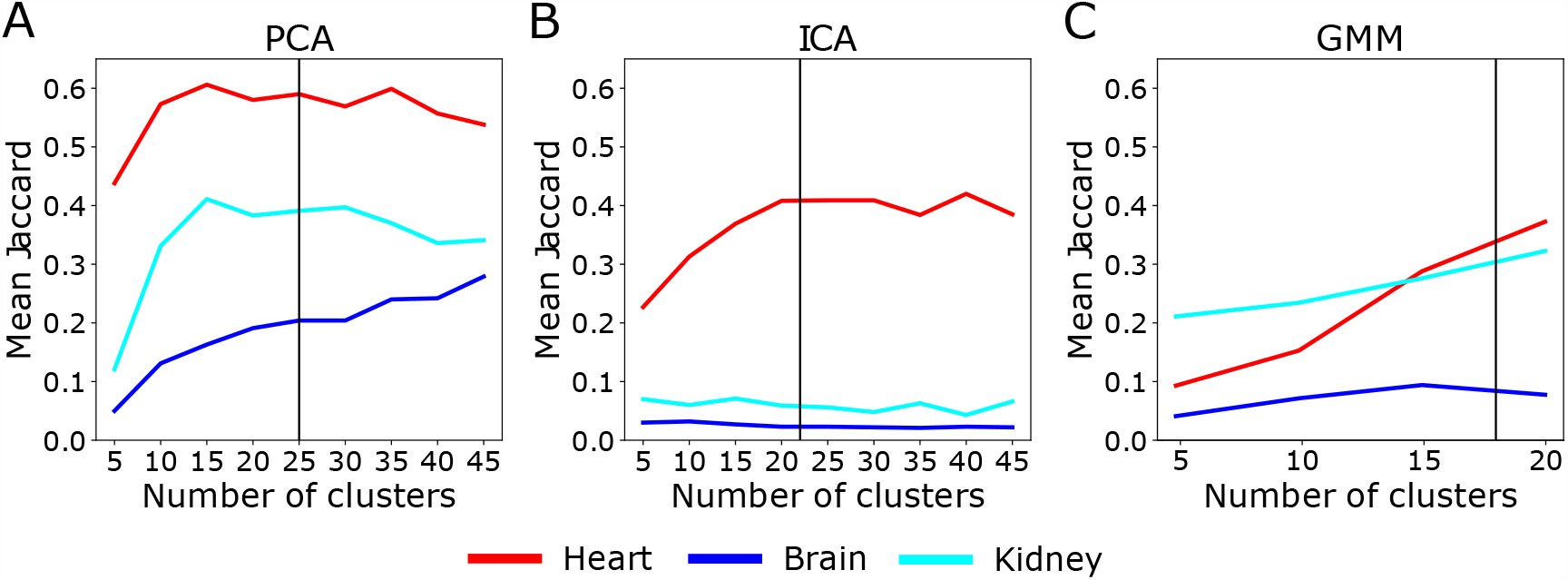
Mean Jaccard index over 10 images (y-axis) as a function of the number of clusters (x-axis) for (A) k-means with principal component analysis (PCA), (B) k-means with independent component analysis (ICA), and (C) Gaussian mixture models (GMM). The accuracy represented by mean Jaccard index is calculated separately for heart (red lines), brain (blue lines), and kidney (cyan lines). The selected number of clusters for further analyses is indicated with a black vertical line.

### 4.2. Clustering results

Next, the remaining 30 images were clustered using the method specific cluster numbers defined in the previous step. ICA combined with k-means failed systematically to identify brain and kidney, and also with the VOI representing heart, its performance was poorer than those of PCA and GMM (Figure 3A). All Jaccard indices are available in numeric format at https://github.com/rklen/Dynamic_FDG_PET_clustering. The performance of GMM comes closer to PCA than expected based on the 10 images analysed earlier as the mean Jaccard indices differed no more than 0.08 units in any of the organs (Figure 3B). Notably, in the context of the heart, PCA has steady performance and GMM has higher variation over the images, whereas in the context of the kidneys, GMM has steady performance and PCA has considerable variation in its performance (Figure 3A and Figure 3C). The differences between the methods were statistically significant in most organs, except in comparisons PCA vs GMM for kidneys and ICA vs GMM for heart (Table 2).

**Table 2.** P-values from Wilcoxon’s test for all organs (columns) and method comparisons (rows) rounded to three decimals.

|  | heart | brain | kidneys |
| --- | --- | --- | --- |
| PCA vs ICA | 0.002 | 0.000 | 0.000 |
| PCA vs GMM | 0.038 | 0.000 | 0.064 |
| ICA vs GMM | 0.339 | 0.000 | 0.000 |

**Figure 3.**
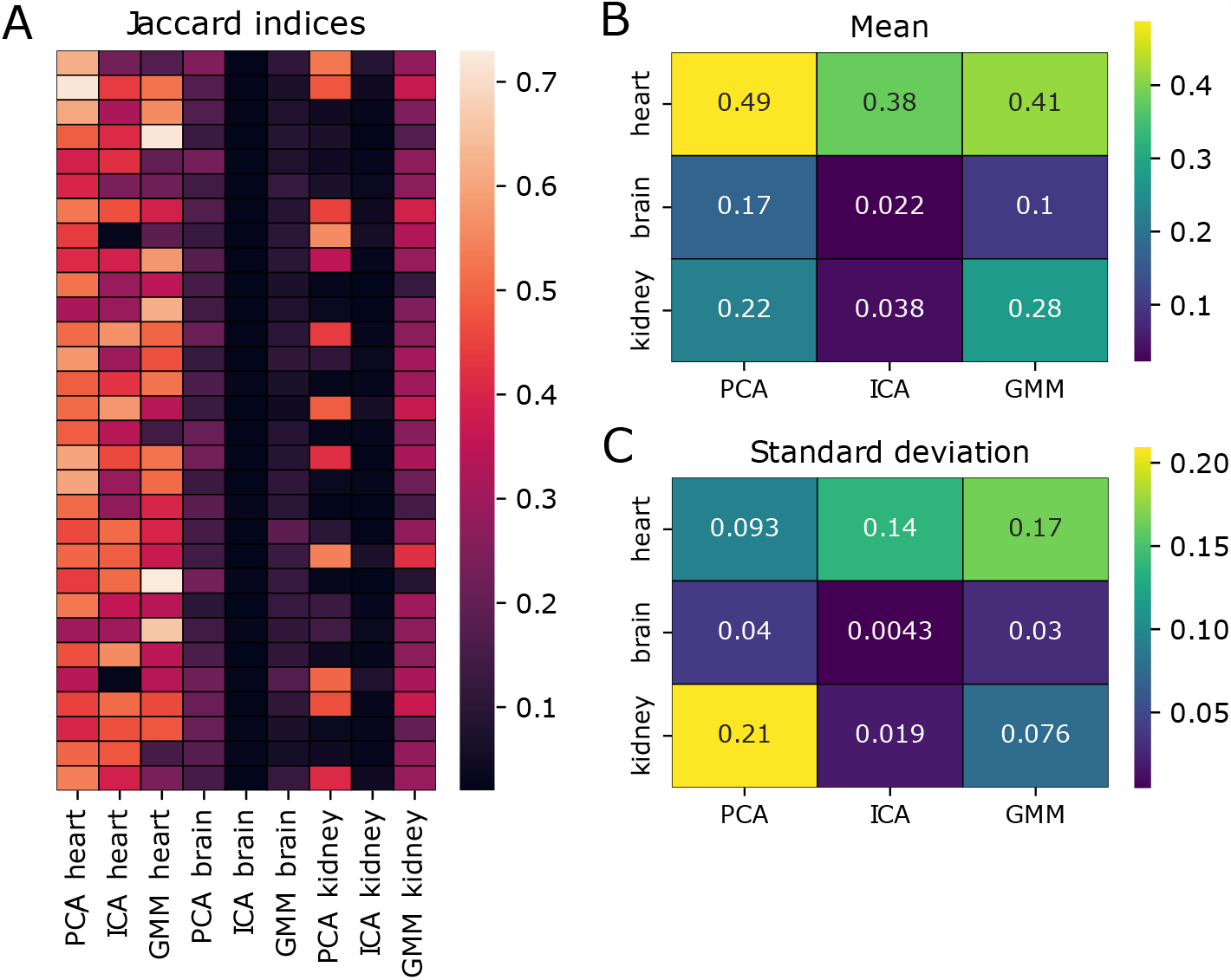
(A) Jaccard indices encoded by colour for different organs and methods (cols) in each 30 test image (rows). The number of clusters was 25 for PCA, 22 for ICA, and 18 for GMM. Subfigures (B) and (C) indicate the mean and standard deviation of the Jaccard indices over the images for all organs (rows) and methods (columns).

Figure 4 illustrates the manually segmented VOIs together with clusters representing them from all the tested methods in one example image. The results for brain are poor because it is clustered with wide areas all over the rat with all the tested methods. The same observation can be made from all the images (Supplementary Figure 3). Interestingly, all of the three methods systematically cluster the spinal cord together with the brain (Supplementary Figure 3), which is reasonable considering their functional connection with nervous system. Other intriguing observations common for all the tested methods are that 1) the heart tends to cluster together with harderian glands (heart-shaped areas behind eyes, Figure 4, Supplementary Figure 4), and 2) brain cluster often includes also brown fat in neck area (Figure 4, Supplementary Figure 3). High radioactivity of harderian glands and brown fat with [^18^F]-FDG has been reported in the literature [43, 44]. GMM is less prone to cluster heart with the harderian glands than the two k-means approaches, but this kind of clustering occurred also with GMM in about half of the test images (Supplementary Figure 4). In some cases, kidneys or heart consist of multiple clusters not spanning over other areas (Supplementary Figure 5). This phenomenon lowers the Jaccard indices artificially in those cases, yet the identified sub VOIs with distinct TAC profiles can be clinically interesting findings, though hard to validate. See section Discussion for further debate about the limitations of the validation.

**Figure 4.**
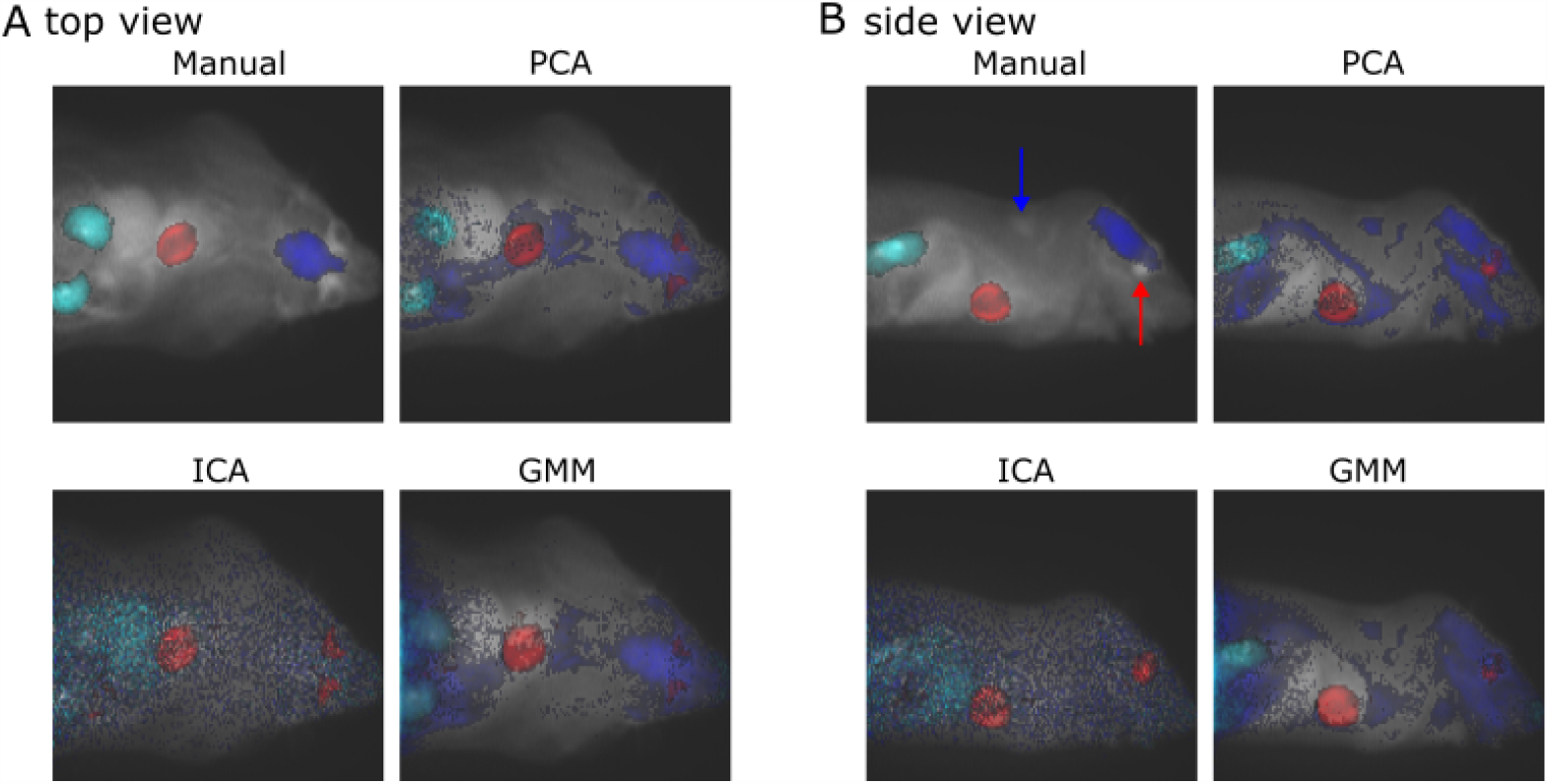
An example image with VOIs identified either manually (topleft panel), or using PCA (top-right panel), ICA (bottom-left panel) or GMM (bottom-right panel). The image is flattened into 2D from (A) top or (B) side view by taking the sum of voxel values over the dimension not visible in the 2D representation. The underlying gray scale image is the PET image and the VOIs are plotted on the top with VOIs corresponding to heart, brain and kidneys encoded with red, blue, and cyan, respectively. The harderian glands are highlighted with a red arrow and brown fat with a blue arrow from the side view image (B) with the manual segmentation.

PCA failed to segment kidneys in 18 test images (Jaccard *<* 0.15). In some instances, kidneys or parts of them were clustered with heart, as illustrated in Figure 5 together with a successful example. Another common issue was kidneys clustered together with larger abdomen area including intestines (Supplementary Figure 6). Supplementary Figure 7 shows that PCA detected kidneys more successfully from the images, where the kidneys fitted better into the scanned part of a body.

**Figure 5.**
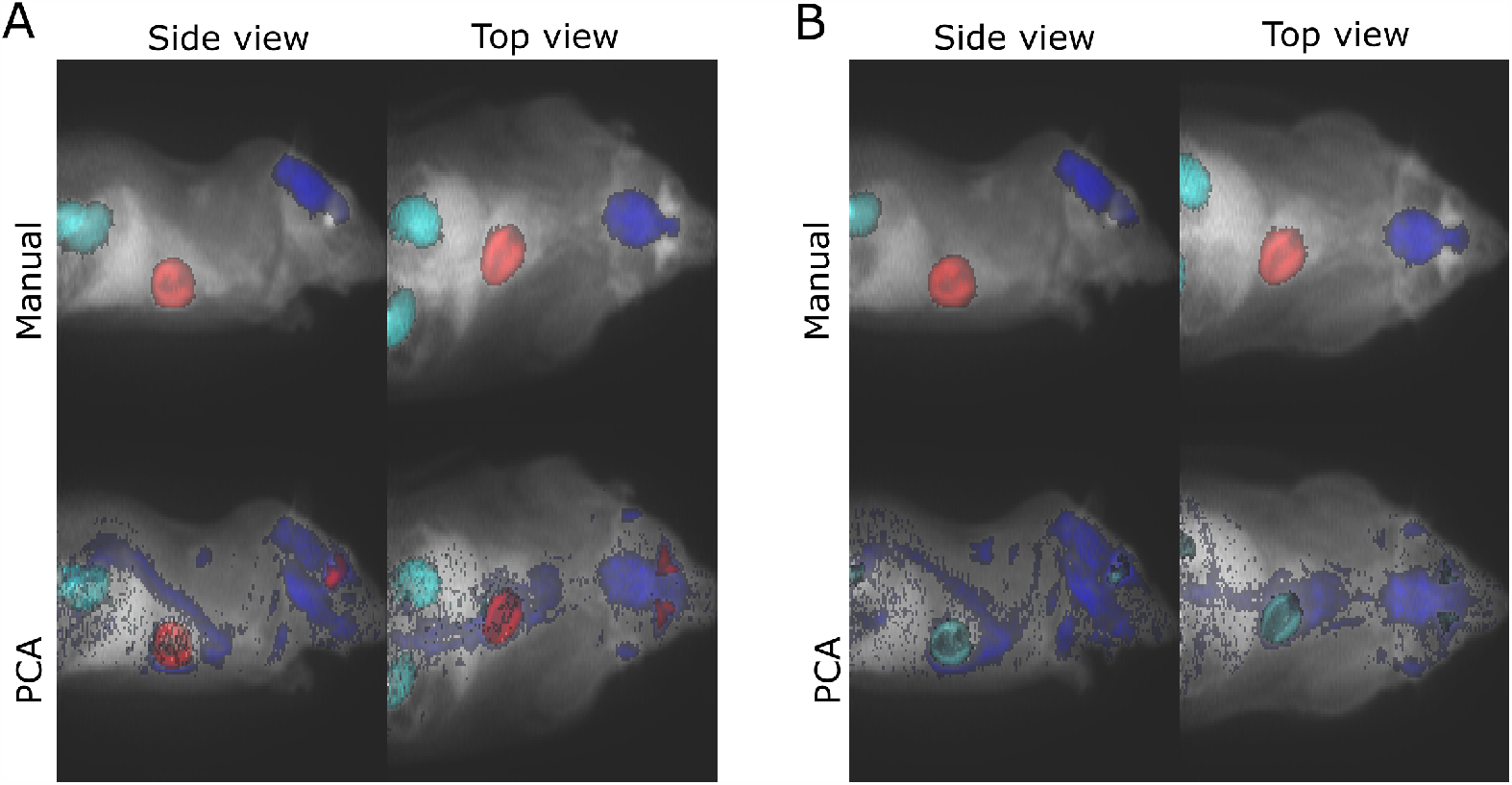
In image (A) PCA succeeded to identify kidneys (Jaccard 0.54), whereas in image (B) it failed in it with Jaccard index 0.05. In both subfigures, the two upper panels include manual segmentations of the VOIs (red for heart, blue for brain, and cyan for kidneys), and the bottom panels visualise the PCA clusters representing corresponding VOIs (again, red for heart, blue for brain, and cyan for kidneys). The underlying gray scale image is the PET image. For visualisation purposes, the images were flattened into 2D format by taking the sum over the dimension missing from the 2D representation.

## 4.3. Human images

We also tested the three clustering approaches on human images. While the sample size of three dynamic 3D images prevents any statistical analysis, these examples are in-line with our observations from the rat data. PCA appears as the most accurate approach, while ICA provides the least intuitive clustering (Figure 6, Supplementary Figure 8).

**Figure 6.**
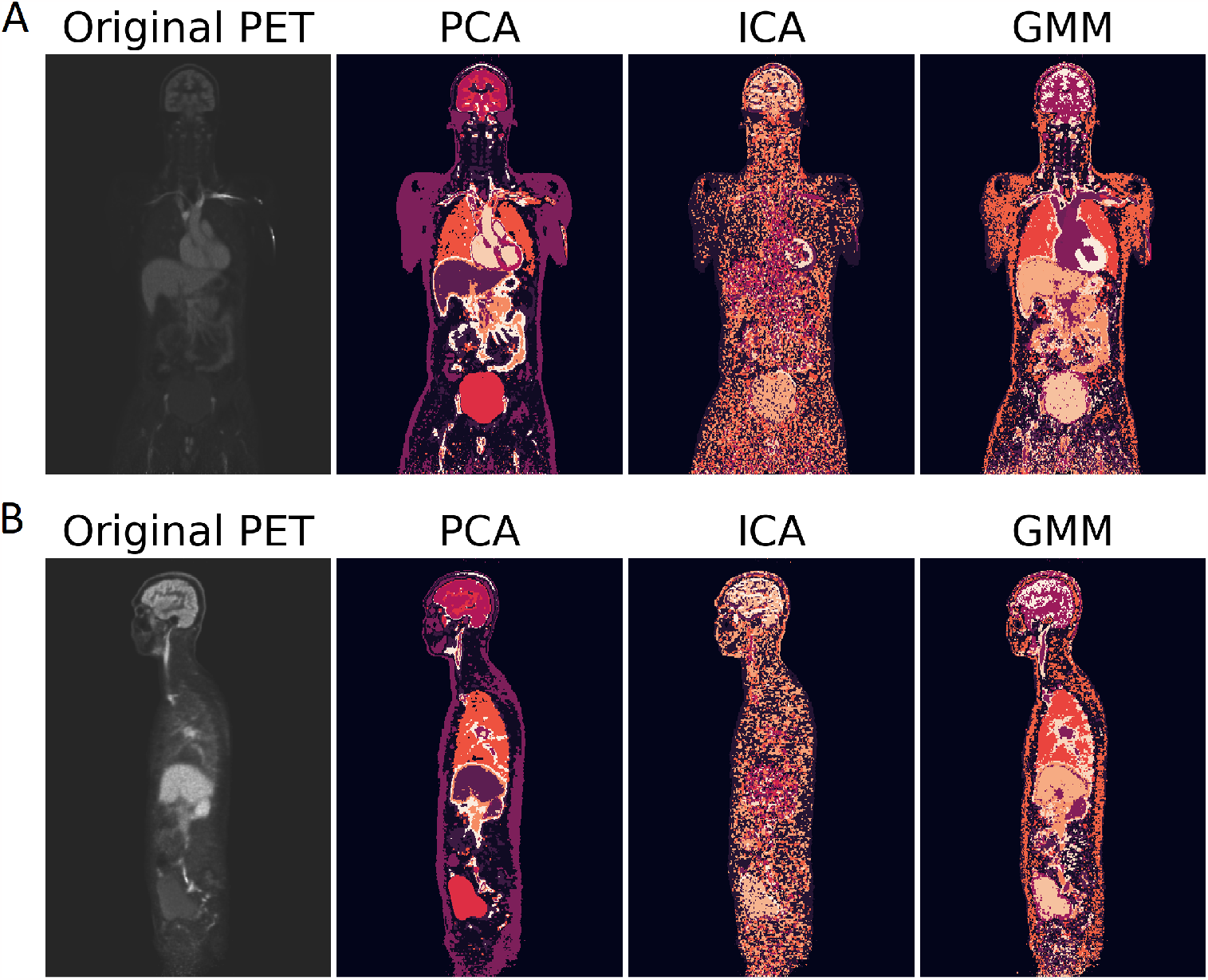
(A) Front and (B) side view of the clusters from the middle of a dynamic 3D human total-body PET image. The visualised slice in both subfigures is the slice number 200 out of 400. Notably, in the original PET image (first panel), the voxel activities are sums over time and brighter colour indicate higher activity, whereas in the cluster images (the remaining three right-most panels) different colours indicate different clusters in no particular order.

## Processing time

Among the tested methods, GMM was 23 times slower than the fastest approach ICA with a mean processing time of approximately 22 minutes per image. Unsurprisingly, the k-mean based approaches were more similar to each other with mean clustering times of 79 seconds (PCA) and 57 seconds (ICA). The standard deviations of the running times for PCA, ICA, and GMM were 14 seconds, 7 seconds, and 5 minutes, respectively. The full list of processing times is available at https://github.com/rklen/Dynamic_FDG_PET_clustering. Notably, with the three human images, PCA was faster than ICA and GMM was only about 10-fold slower than the k-means approaches. The mean running times (rounded to full minutes) with human images for PCA, ICA, and GMM where 7 minutes, 10 minutes, and 73 minutes, respectively.

## 5. Discussion

The ultimate purpose of studying the clustering of PET images is to aid the process of quantification of PET data. In the end, the clustering could be used to automatically segment PET images into VOIs that could be used in kinetic modelling. This is an early investigation on clustering total-body images and many new applications are still possible. This kind of total-body clustering could possibly find connections between TACs of different tissue types, which could give insight on different problems in medicine. For example, total-body clustering could be experimented in finding metastatic tumors and otherwise just different tissues that have the same uptake of the tracer. Organ segmentation also supports diverse system level applications of image analysis [45, 46, 47, 48].

We suspect that among the tested VOIs, heart is the easiest one to segment with clustering due to the high and relatively homogeneous maximum activity values of its TACs (Supplementary Figure 2). Similarly, among the tested VOIs, the brain was the most difficult to detect with clustering and it has functionally heterogeneous areas and lower radioactivities than the other two organs. This suggests that the overall activities have an important role in the clustering process. Notably, PCA had unstable performance with kidneys depending on how well the kidneys fitted into the scanned area, but it is unclear why only PCA was sensitive to this. However, it is also probable that whole organs may not fully fit into human total-body scans. In this study, we used PET images containing [^18^F]-FDG tracer, which is not commonly combined with dynamic imaging in clinical usage. However, for our technical evaluation this is not a problem, as clustering should not be sensitive to the selected tracer as long as the manual segmentation used as gold standard is drawn from the same images used for clustering.

Notably, the original motivation to use GMM arises from the voxels of a VOI in a static 3D PET image typically having a distribution easily modelled with a mixture of Gaussian distribution. For dynamic images where the distributions are fitted into TACs this is not necessarily the case. Thus, GMM might be better suited for segmenting static 3D PET images than dynamic 4D PET images.

Due to the differences in the scanned area and evaluation metrics, it is difficult to compare our results to the results obtained in the previous studies. However, higher accuracies have been achieved in the brain region level with sophisticated yet computationally demanding methods [8, 49]. This highlights the need for a better clustering approach designed for very large images.

Different machine learning approaches are fast to use after the training has been done, which make them an appealing approach for segmenting very large total-body images, though the available training data is still very limited. However, the scope of their segmentation is different from that of in-situ approaches like clustering. Learning-based machine learning methods can be used to segment VOIs that are expected to be present in all images (such as full organs), whereas in-situ approaches are suited to segment VOIs that differ from the rest of the image in the particular image under study (e.g. part of liver behaving unexpectedly). The research question defines which type of segmentation is desirable.

This is an early stage study with several limitations in the test data and evaluated methods. Besides the clustering approaches being used as such without any further processing discussed here, the only tracer utilised was [^18^F]-FDG. While it is a widely used one, other tracers, particularly those not based on fluorine, could provide drastically different data as they accumulate differently. We do not expect different clustering methods to be accurate with different tracers, but another tracer could introduce different challenges not present in this study. In addition, our data was limited to 40 images from rats and 3 images from humans, while ideally a larger number of human total-body images would have been used. For the sake of straightforward validation, only three VOIs were considered here. In real application, the number of functionally distinct areas of total-body PET images is much greater, though most studies likely focus on only some of them. Other issue with the validation is that the manual segmentation is done on sum 3D images neglecting the time aspect. Thus, VOIs with different time-patterns, but similar total radioactivities won’t be distinguished and segmentation methods for static 3D PET images would likely appear equally strong to the methods for dynamic images if compared here. However, doing the manual segmentation for validation purposes from a video of a 3D object is not feasible at the time of writing.

Notably, bladder filling during the imaging process has been stated to be a problem for PCA in the literature [18], but our data does not support that claim. While the bladder is not within the scanned area in our rat data, the human images include it, yet PCA provided meaningful clustering on them. However, our human data is limited to three images so further investigation of the topic is needed.

## 6. Conclusions and future work

Here we have tested three clustering approaches to identify predefined VOIs from dynamic total-body PET images based on voxel specific TACs. Data from rats and humans were used, but the specie did not have high impact on our conclusions. Our results show that none of the tested methods is accurate enough for practical applications without further method development, but at least the tested methods were still computationally usable also with the large human images. The obtained mean Jaccard indices reflecting the performance of the tested methods varied from almost 0 to 0.17 for the most difficult organ brain, from almost 0 to 0.28 for the kidneys, and from 0.38 to 0.49 for the heart. Independent component analysis provided the weakest results with all organs.

One possible approach to prevent entirely unrelated parts from clustering together is to split the images into smaller subsections or otherwise limit the clustering to nearby voxels. Alternatively, splitting the unconnected clusters into separate ones could be automatically done with connected component analysis as a post-processing step. Besides preventing scattered clusters, this would also allow using slightly more computationally demanding methods and, in the case of dividing the image into subimages, different number of clusters could be used in different image regions. The last benefit seems important, as in this study, the brain benefited from larger number of clusters than heart or kidney when PCA approach was used. Particularly complex and heterogeneous VOIs like brain could benefit from this approach. The cons of looking only for connected clusters include, for example, difficulties with associating metastases with the primary tumour, and pair organs like kidneys not clustering together. A post-processing step where tiny (particularly single voxel) unconnected parts of a bigger cluster would be removed from their original cluster and merged into the attached cluster that most resembles them could be a reasonable compromise with little to loose between the pros and cons of looking for connected clusters. We had to exclude majority of the previously used approaches for computational reasons, which highlights the need for computationally lighter solutions. These can be obtained by implementing new computationally light segmentation methods, implementing work-arounds that allow the use of current methods (e.g. splitting the images into smaller pieces as suggested above), or different hardware and cloud solutions. For example, service-oriented networks have been shown to reduce the computational cost of image analysis as compared to corresponding software [50].

Other interesting topics for further research include the automatic definition of number of clusters, the possibility to combine nearly located clusters if they resemble each other, pre-processing of the imaging data, and the use of kinetic models. The optimal number of clusters is likely heavily dependent on the size of the image and the utilised tracer, so we do not expect that the cluster number defined in this study generalises well to other studies. Different pre-processing approaches have the advantage of typically being computationally lighter than sophisticated clustering methods. For example, noise reduction by using mean activities over the voxel neighbourhood instead of the measured activities of a voxel as such could improve the clustering accuracy particularly for ICA, which performed poorly with brain and kidneys, possibly due to being sensitive to noise. Also scaling the TACs so that the clustering is done based on their shapes rather than absolute activity levels is an interesting option already applied in the literature [15, 13]. It could affect which VOIs are easy and which ones difficult to detect with clustering. Also considering issues common to PET data, such as spill-over effect from closely located organs and partial volume effect, could improve the results.

For automatic segmentation to become routinely used in clinical work, implementations of the available tools have to be a) available, b) easy and intuitive to use, and c) compatible with the readily used visualisation/analysis softwares. The first criteria sounds obvious, but particularly the implementations of the old unsupervised methods are often not available. As many clinicians are not experienced programmers, we would like to highlight the importance of the ease of use of the future methods (criterion b and c). While a Python package or other implementation that can be used freely in basically any computer is a must, we would encourage the method developers to provide also an implementation that can be used with the locally utilised visualisation/analysis software like Carimas [42], 3D Slicer [51], PMOD, or AMIDE [52]. This would increase the likelihood that the cutting edge methodology is used also in practical clinical work.

In this study, we have shown PCA combined with k-means to be the best performing method among the tested alternatives. In addition, its running time is in the same scale as the fastest method tested here. However, considering its incapability to capture brain from the images and unstable performance with kidney, there is room and need for a more sophisticated total-body TAC clustering method. While the requirement for scalability to very large datasets presents some challenges, we have discussed multiple computationally light approaches to further improve the results.

## Supplementary material

Supplementary file SupplementaryFigures.pdf includes all the supplementary figures that were referred to in this manuscript.

