## SupplementaryFigures for "SEGMENTATION OF DYNAMIC TOTAL-BODY [^18^F]-FDG PET IMAGES USING UNSUPERVISED CLUSTERING"

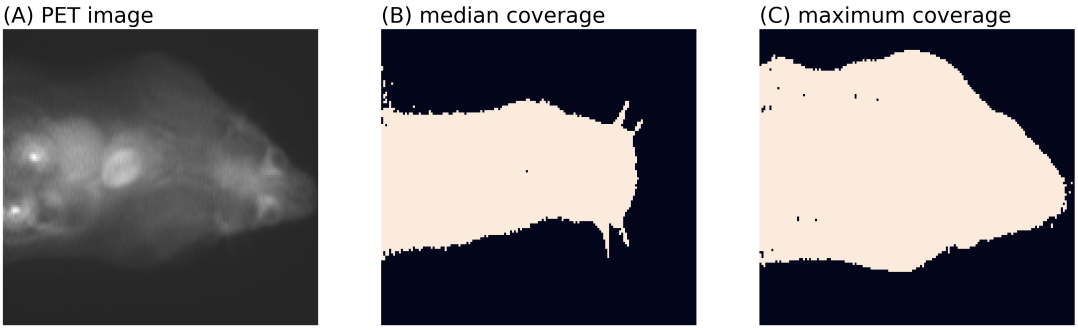


Supplementary Figure 1: (A) The original PET image (top view, sum over time points and the missing dimension) and the voxels of it used for clustering from (B) the slice with median number of clustered voxels (slice 49/86), and (C) the slice with the maximum number of clustered voxels (slice 74/86). The slice with the minimum number of voxels used for clustering is not visualised here, because it would be blank.

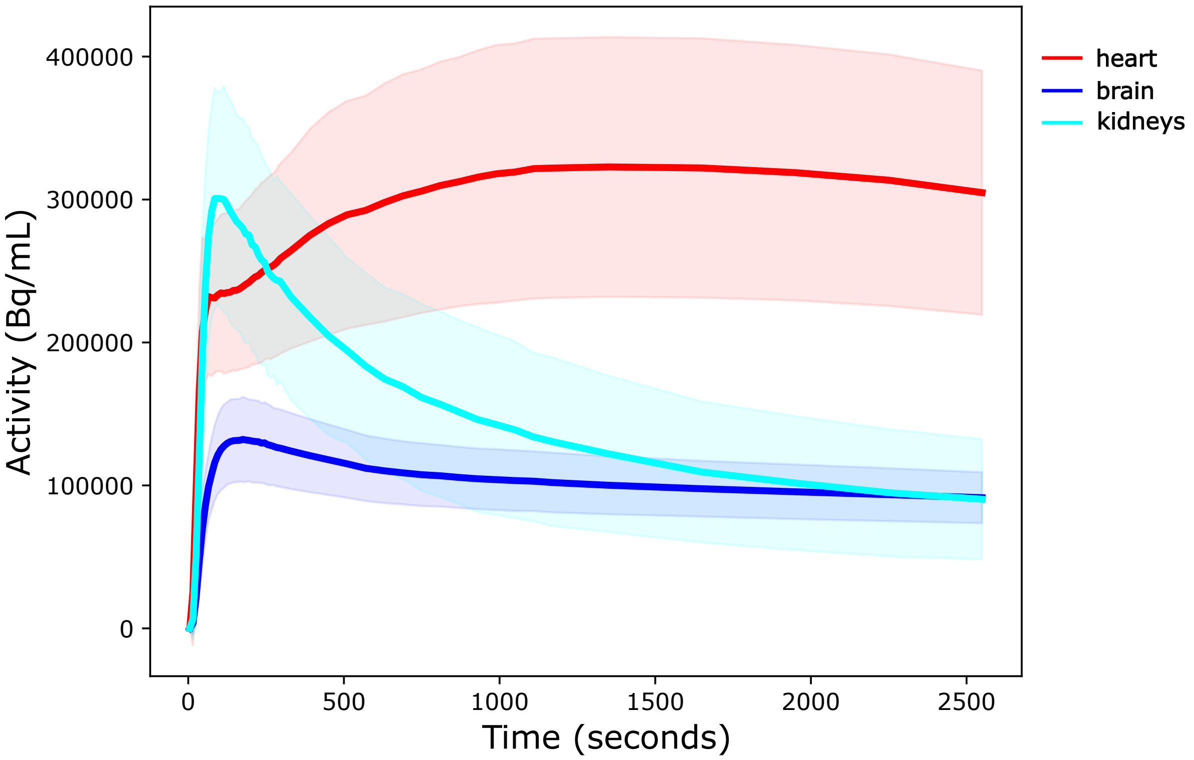


Supplementary Figure 2: Mean time activity curves of heart (red), brain (blue), and kidneys (cyan). First mean over voxels belonging to each organ was calculated from each image, and then mean of those image specific curves was drawn (visualised here). The semi-transparent areas around the curves indicate the standard deviations over images.

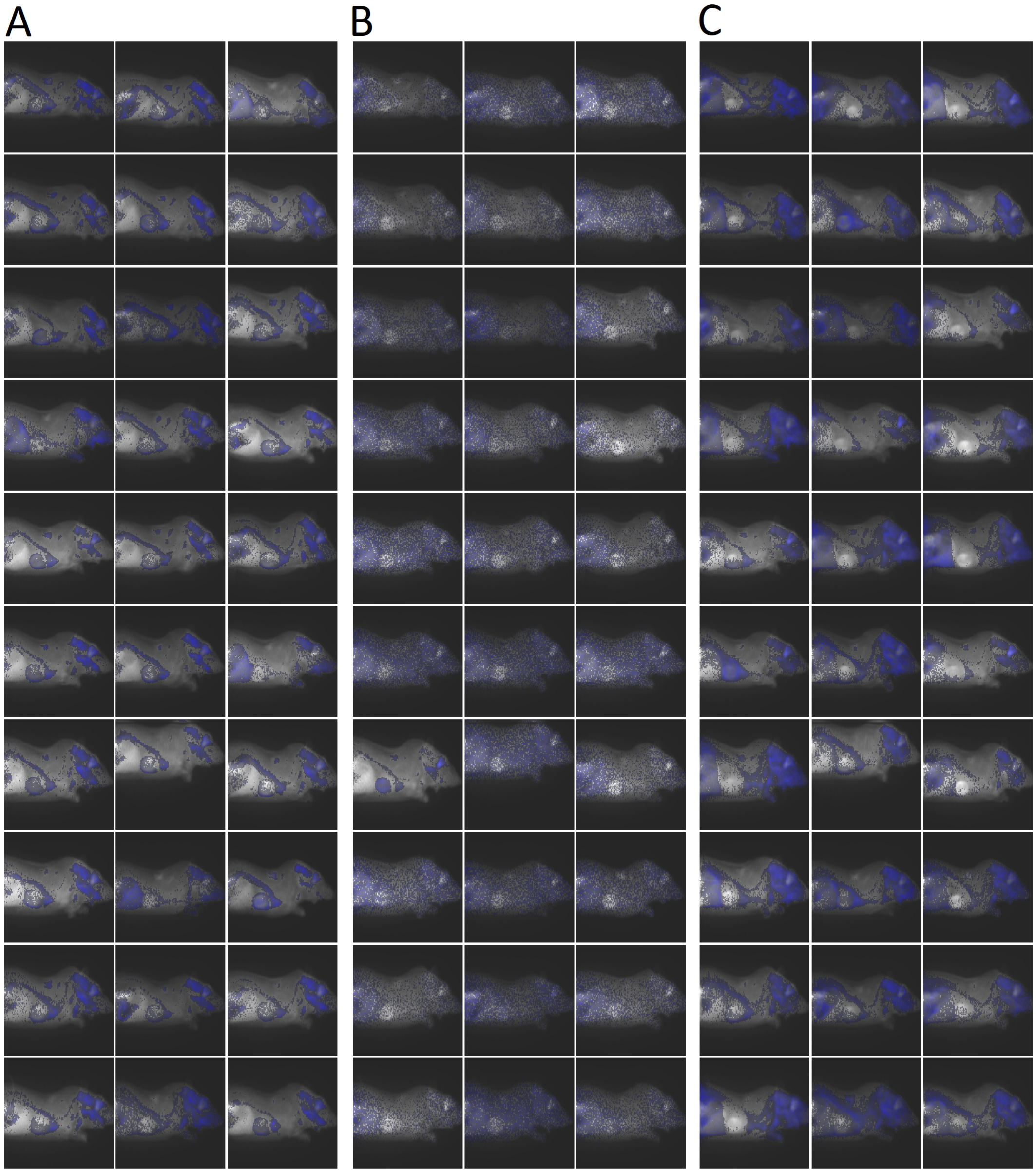


Supplementary Figure 3: Brain clusters for the all 30 test images detected by (A) PCA, (B) ICA, and (C) GMM highlighted with blue colour. The underlying gray scale images are the corresponding PET images for reference.

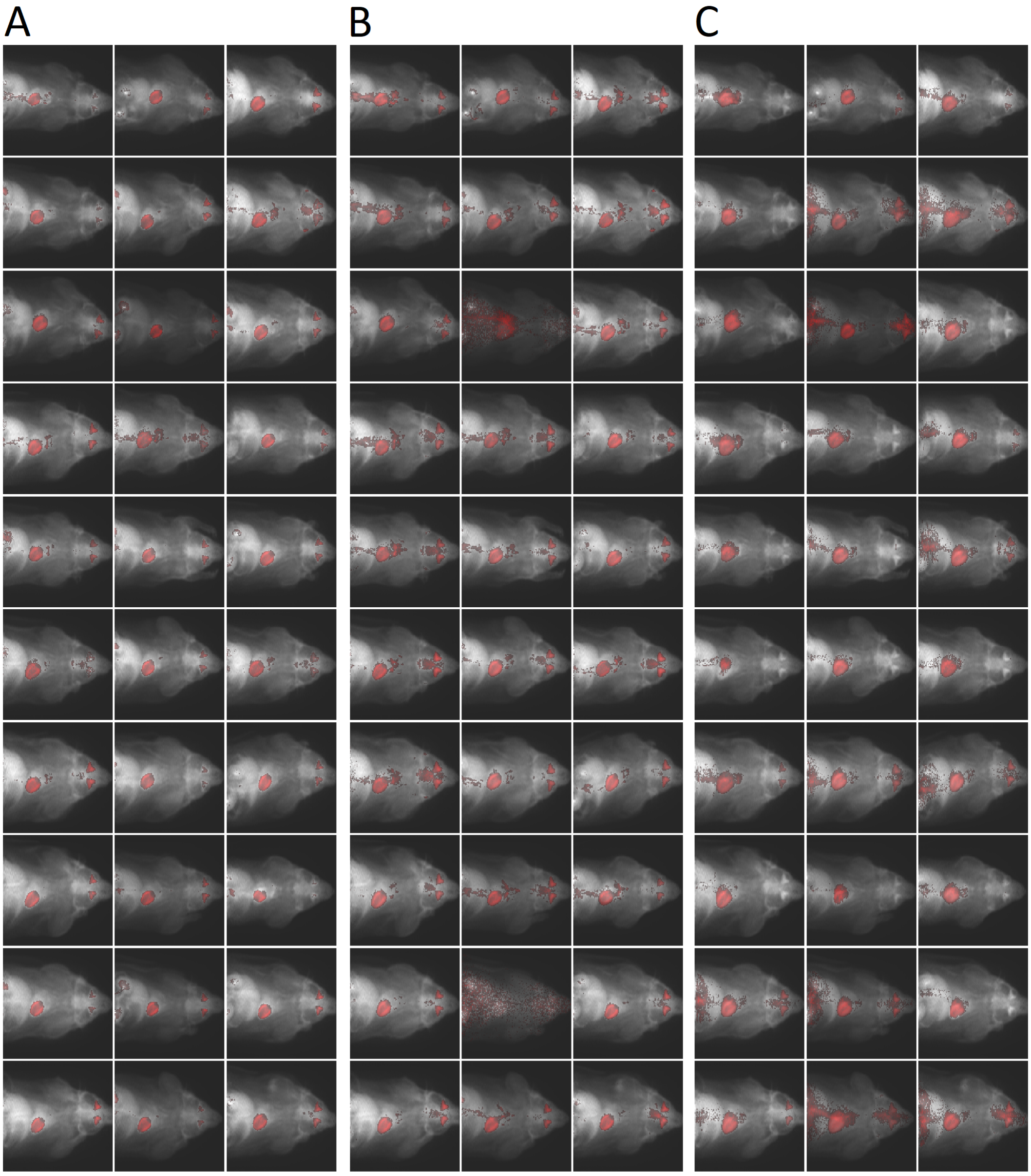


Supplementary Figure 4: Heart clusters for the all 30 test images detected by (A) PCA, (B) ICA, and (C) GMM highlighted with red colour. The underlying gray scale images are the corresponding PET images for reference.

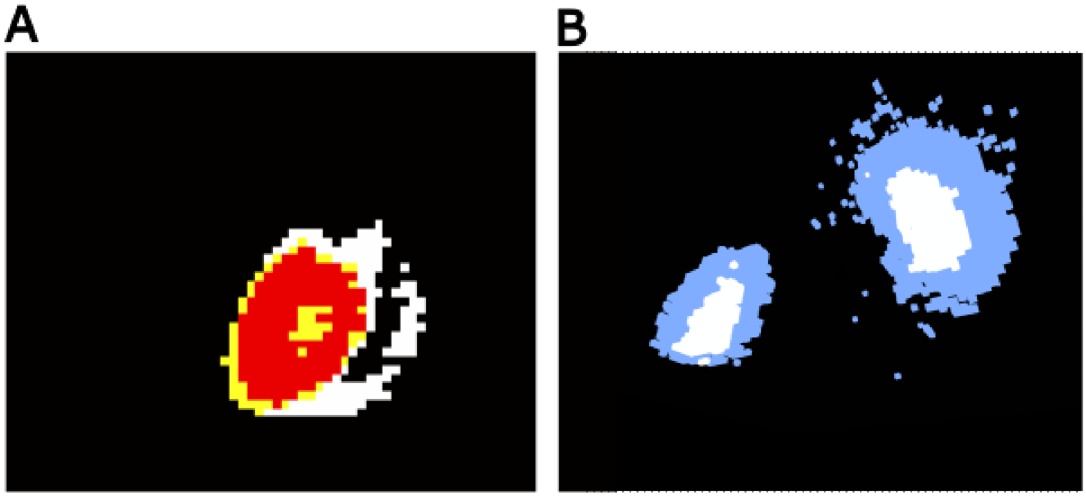


Supplementary Figure 5: (A) Heart and (B) kidneys containing 3 and 2 clusters, respectively. The heart and the kidneys are from different PET images.

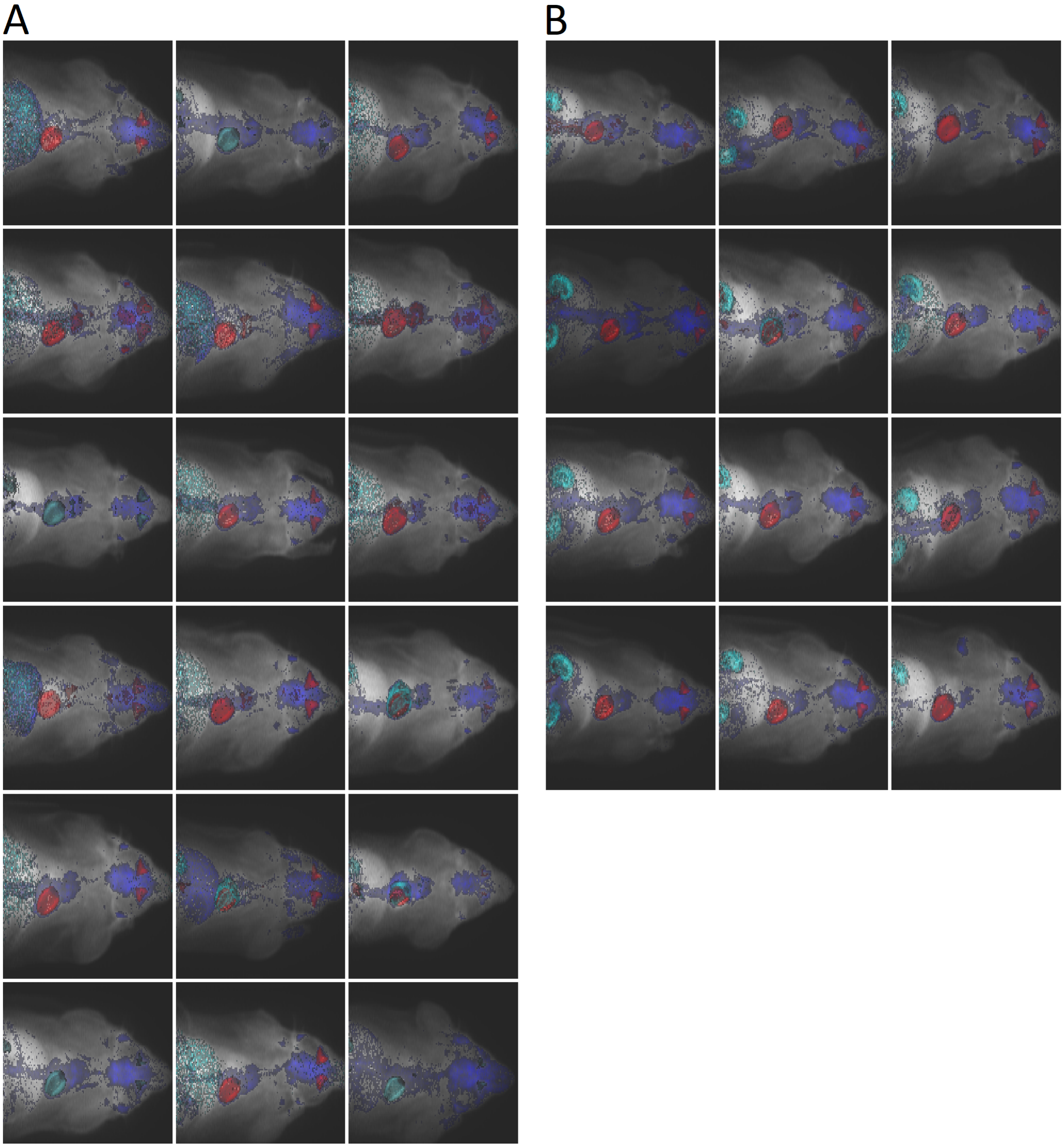


Supplementary Figure 6: Instances where PCA (A) failed to identify kidneys and where it (B) succeeded at it. PCA clusters for heart, brain, and kidneys are indicated with red, blue and cyan colour, respectively. The underlying gray scale images are the PET images for reference.

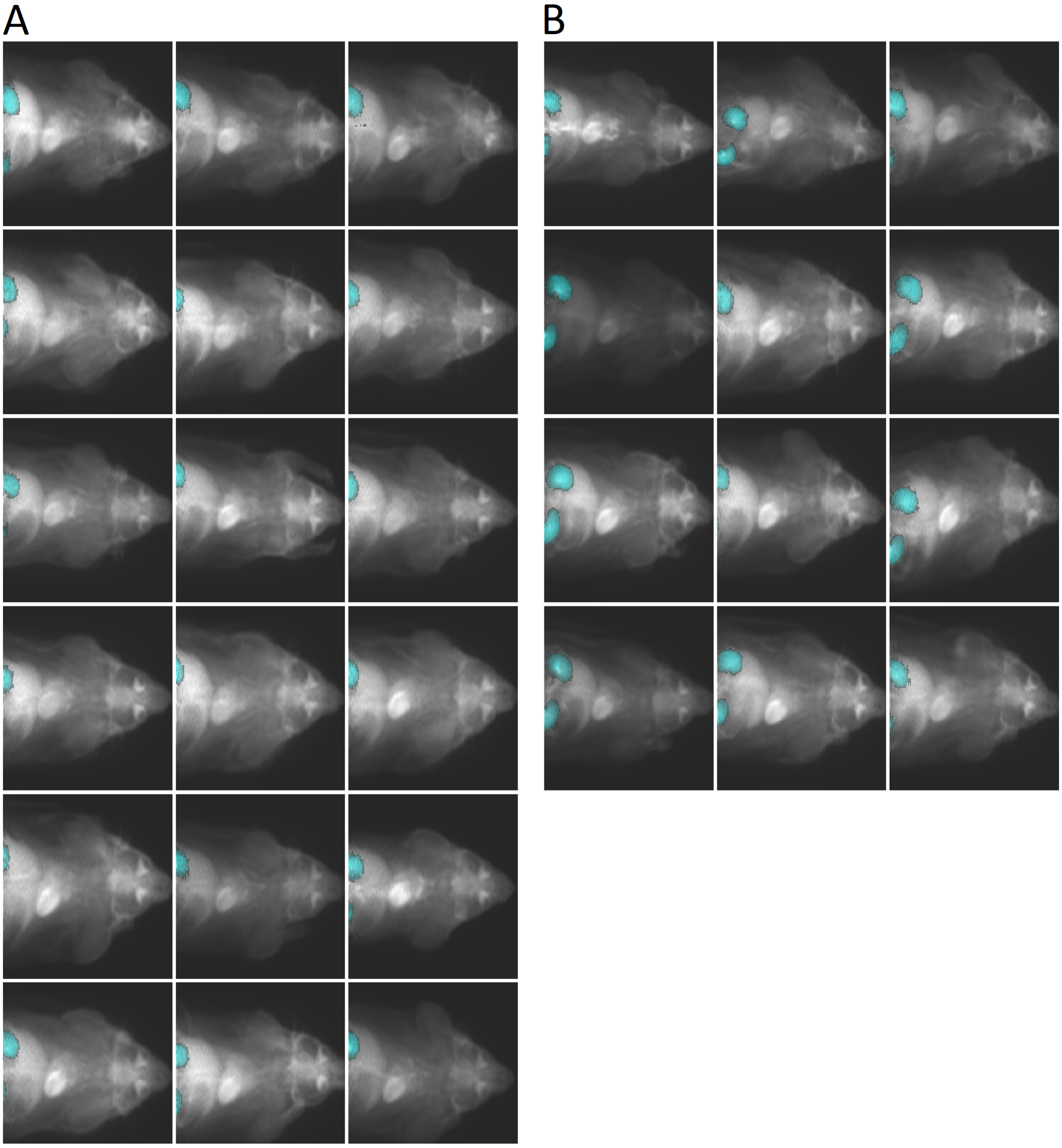


Supplementary Figure 7: Instances where PCA (A) failed to identify kidneys and where it (B) succeeded at it. Manually segmented ROIs for kidneys are indicated with cyan colour. The underlying gray scale images are the PET images for reference.

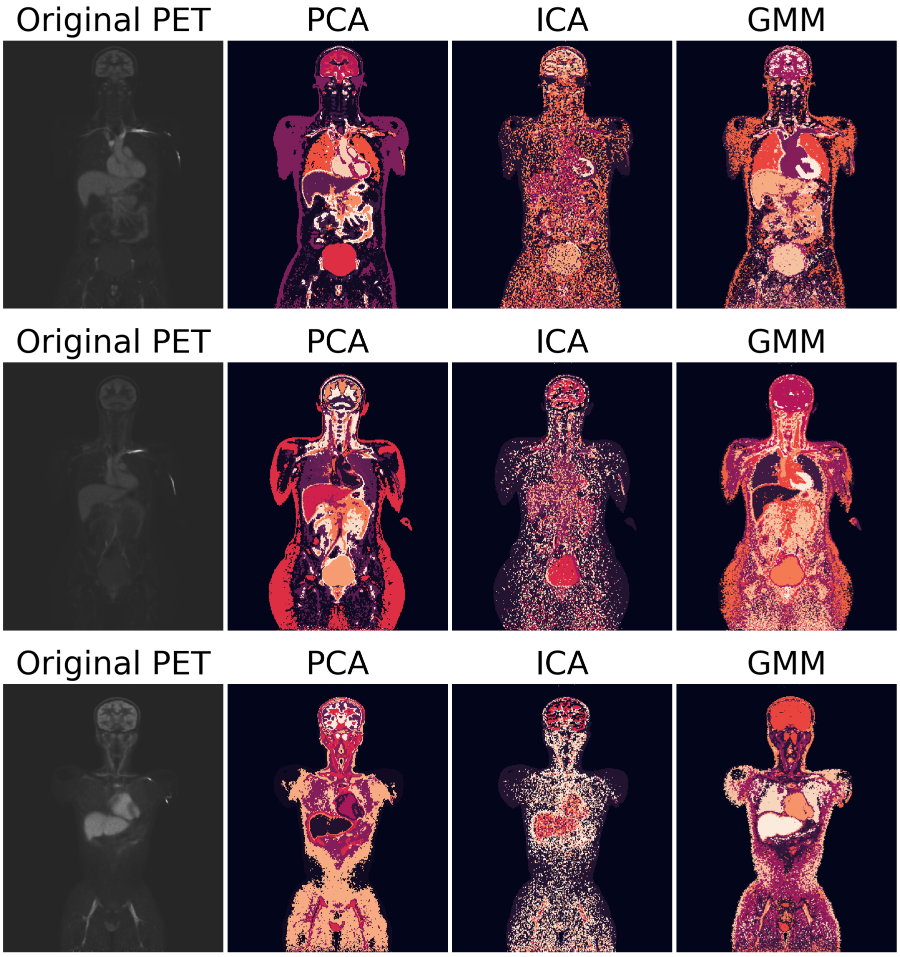


Supplementary Figure 8: Clusters defined using different methods (columns) for all three dynamic human total-body PET images (rows). The visualised slice is the slice number 200 out of 400. The gray-scale images on the left-hand side are the corresponding slices of the original PET images, where the colour reflects the total activity over time in each voxel of the slice.
